# Does winterkill explain contrasting demographics of winter-breeding freshwater mussel populations in Ishikari River floodplain?

**DOI:** 10.1101/2023.11.07.565987

**Authors:** JN Negishi, H Izumi, J Wu, S Fukui, I Koizumi

## Abstract

1. Environmental conditions bottlenecking species population demographics are less known in cold regions with harsh winters despite of global concerns of declining freshwater mussels (Unionidae). Few studies examined both cold and summer environments in attempts to promote habitat conservation of Unionidae species.
2. We identified the taxonomically confused Unionidae species using phylogenetic analysis with mtDNA (COI region) and tested the hypothesis that winter mortality is the main cause of contrasting population demographics and structures in floodplain lakes in northern Japan. Demographic surveys were conducted in two lakes, which contrasted in recruitment rates, over one year including 4-month ice-covered periods
3. Although previous studies have identified freshwater mussels as introduced *Anemina arcaeformis* (Heude 1877) based on morphology, this study confirmed the focal species as potentially native *Buldowskia iwakawai* (Suzuki, 1939). High winter mortality (30-40%) of adult mussels was found, although the mortality did not significantly differ between the populations with contrasting recruitment.
4. Surprisingly, the annual mortality was much lower in juveniles (10%) than in gravid and nongravid adult individuals (40-75%). The main inter-population difference was attributed to the higher summer mortality of gravid females, but not juveniles and non-gravid individuals, in the population with low recruitment.
5. These results collectively suggest that summer hypoxia combined with physiological stresses on females in winter is a likely population growth-limiting mechanism. To prevent a chain of adult abundance decreases in winter and high mortality of gravid mussels and newly born juveniles in summer, improvements in summer habitat conditions are necessary, while winter conditions need to be considered simultaneously. Increases in water circulation rates and alleviations of hypoxic conditions is an option for short-term habitat improvement approach. The current study sheds light on the winter-mediated mortality of freshwater mussels in shallow eutrophic floodplain lakes and contributes to improved management strategies for degraded floodplain waterbodies.

## Introduction

Floodplain waterbodies are among the most threatened ecosystems on Earth because of multiple anthropogenic stressors, including land use change, water pollution, and hydrological cycle disruptions (Tockner & Stanford, 2002). The ecosystem management challenges of multifunctional floodplain waterbodies concern the maintenance of their ecosystem services, one of which is the provisioning of habitats for diverse organisms against increasing human demands and the development of floodplain areas because of easy access to water, fertile soils, and flat land readily used for various purposes while preventing floods (Schindler et al., 2013, 2016). Alterations in the flow regime and hydrological connectivity in floodplain-river ecosystems have received much attention as key considerations in the conservation and restoration of degrading floodplain habitats and biodiversity worldwide (Bunn & Arthington, 2002; Poff & Zimmerman, 2010; Górski et al., 2012; Rabuffetti et al., 2017; Melack & Coe, 2021). Current findings on floodplain ecosystems are biased towards ice-free seasons due to year-round warm climates or lack of such efforts in ice-covered seasons although degradation of floodplains is pervasive regardless of the underlying climatic conditions. The winter period can potentially be a bottleneck in the distribution of organisms in relatively productive floodplain lakes, as exemplified by winter hypoxia and kills of animals, that is, winterkill, which has long been known for non-floodplain lakes (e.g., Magnusonl et al., 1985; Hilt et al., 2014).

Freshwater mussels (Unionoda) are filter-feeding benthic invertebrates and a common group of organisms inhabiting floodplain waterbodies (Colle & Callil, 2012; Negishi et al., 2012; Negishi et al., 2018; Ćmiel et al., 2020). They are known to have important ecological functions (Vaughn et al., 2004; Strayer, 2014). Some populations of freshwater mussels are endangered (Bogan, 1993; LopesLLima et al., 2017; Zieritz et al., 2018), and this trend is partially due to their important habitat degradation in floodplains. Their relatively complicated life cycle is characterized by an obligate parasite stage that requires the host fish for reproduction (Strayer, 2008; Bauer & Wächtler, 2012). Known stressors for freshwater mussels include direct environments, such as temperature rise (Pandolfo et al., 2010), water pollution and quality degradation (Strayer & Malcom, 2012), habitat loss for them and their hosts (Vaughn & Taylor, 1999; Negishi et al., 2013), and indirect causes, such as host fish decrease (Tremblay et al., 2016), and competition with invasive species (Douda et al., 2012). Breeding seasons differ among species, and some species are adapted to breeding in the middle of winter (Kondo, 1989; Roberts & Barnhart, 1999). Winter-breeding species are less studied in general compared to summer species, and our understanding of their ecology in areas with long or severe winters and field surveys is highly limited in a few cases (Negishi et al., 2018), despite the need to search for a cause of population decline (Izumi et al., 2020).

One of the most prevalent signs of population decline in freshwater mussels is the cessation of their reproduction, which can often be reflected in disproportionately fewer juveniles in population demography and small population sizes (i.e., abundance) (Österling et al., 2010; Negishi & Kayaba, 2010). Efforts have been made to identify how population decline and reproduction cessation occur by examining single or multiple life-history stages of species in relation to various external environmental factors (Geist et al., 2006; Österling et al., 2010; Strayer & Malcom, 2012; Tremblay et al., 2016; Brian et al., 2021; Miura et al., 2023a). Furthermore, as intrinsic properties of mussels, demographics, including size frequency, age structure, sex ratios, and survival rates, are among the key parameters in inferring impaired populations and projecting how and whether the populations would change in the future (Ferreira-Rodríguez et al., 2019). Therefore, field observations of how populations are maintained in different ways in a wide range of environments, from benign to degrading habitats, are needed. Numerous studies have reported the life history and demographics of Unionoida mussel populations (e.g., Negus, 1996; Mcivor & Aldridge, 2007; Jones et al., 2010; Negishi & Kayaba, 2010), but reports on how this taxon varies in multiple aspects such as population-level size distribution, individual-level mortality in relation to their reproductive activities, and life-cycle stages under different environments are still limited (Matter, 2013) and absent in harsh winter contexts.

We previously reported that populations of freshwater mussels, *Anemina arcaeformis* (Heude, 1877), are in trouble with their reproduction, as measured by the proportion of juvenile mussels, which ceased in most floodplain lakes of the Ishikari River (Negishi et al., 2018; Izumi et al., 2020). Freshwater mussel taxonomic classification has recently been substantially revised by a supplementary powerful aid of genetic information and subsequent phylogeny analyses in the East Asin region (Lopes-lima et al., 2020), in which one individual of *Buldowskia iwakawai* (Suzuki, 1939) was reported from the same lowland area of the Ishikari River floodplain. Although these two species are morphologically similar, their ecological status and implications differ because only the latter is native to Japan. As the Ministry of Land, Infrastructure, Transportation and Tourism (MLIT) of Japan currently promotes environmental restoration plans for degrading natural floodplain environments in the area (MLIT, 2014) by identifying causal environments that bottleneck population and distribution increases (survival and reproduction), correct identification of this unionoid species (hereafter Unionidae sp.) and understanding of their distribution is necessary. In this study, we identified the species using phylogenetic analysis with mtDNA (COI region) and tested the hypothesis that winter mortality is the main cause of contrasting population demographics and structures. Pongsivapai et al. (2020) recently reported that the organic matter content in lake sediment has increased over time, which implies that the sediment condition preferred for an increased level of hypoxia, a major driver of winterkill, has intensified. Based on this report, we predicted that winter mortality is higher than mortality during non-winter periods and is disproportionately higher in the impaired population, regardless of gravidity.

## Methods

### Study site

The field study was conducted from December 2015 to November 2022 in 11 floodplain water bodies (FWBs) along the lowland section (approximately 50 km long) of the Ishikari River (catchment area of 14,330 km^2^) in Hokkaido, northern Japan (Fig. 1). The exact locations of these water bodies remain undisclosed for conservation purposes. The study channel segment had an average gradient ranging between 0.2 and 0.7% and an annual average flow rate of 327 m^3^/s. The Ishikari Plain, including the study area, historically formed a vast area of peatland once exceeding 550 km^2^, as a result of ocean intrusion approximately 6,000 years ago, followed by the gradual retreat of the coastline. The national government led a large-scale drainage project (1950–1960s) to convert marshy areas for human use by straightening the tributaries and main channel. Although the area is still underlain by peat with a thickness of 4–5 m, almost all of the peat marsh has been drained and lost (Miyaji & Kohyama, 1997). Consequently, the originally estimated channel length of the Ishikari River (364 km) was reduced to its present length (270 km). The study FWBs were either “eutrophic” or “hypertrophic” (Pongsivapai et al., 2020). The small and shallow water bodies provided a suitable habitat for macrophytes that led to OM deposits, resulting in an increase in the OM and OM to total nitrogen (TN) ratio over time. Monthly mean(±SD) air temperature in the period ranged −6.6(±0.79) to 21.1(±1.16) and minimized and maximized in the month of January and July, respectively (Bibai weather station, Japan Meteorological Agency within seven km). Surface ice formed (up to ca. 20-cm think) in the studied water body in the early December till the early April.

**Figure 1.**
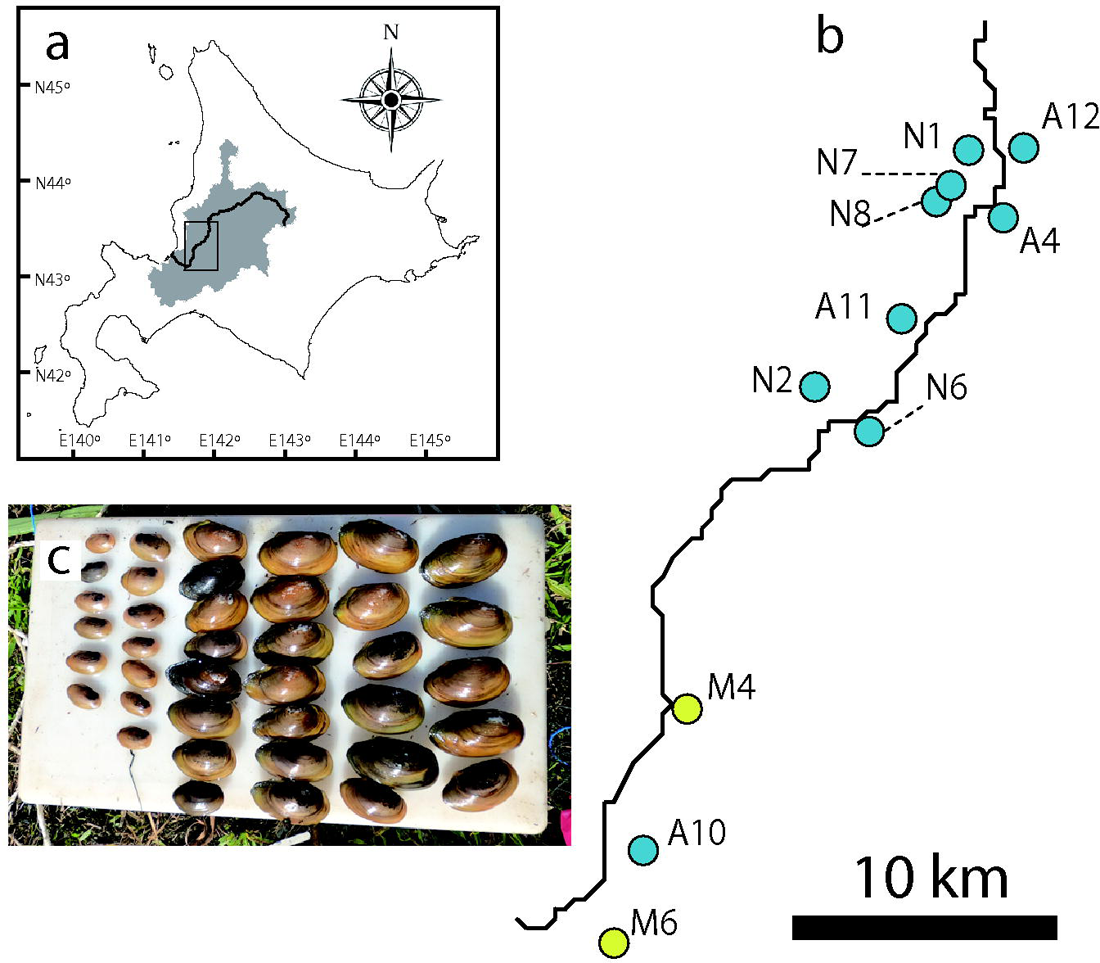
Ishikari River watershed (shown in gray shade) in Hokkaido (a) and the location of study sites in the dotted areas in (a) (b), and a picture showing juvenile unionoid mussels collected including several species (c). In (b), yellow circles indicate study sites where intensive winter monitoring was conducted, whereas all sites, including those shown as blue circles, were used to collect mussel tissue for DNA barcoding analyses.

### Species identification with DNA barcoding

Sites were visited between 2015 and 2022, and Unionidae sp. and sympatric Unionid species were collected (59 individuals; Supplementary information 1). Small pieces of foot tissue were cut and preserved in 99.5% ethanol until further analysis. Taxonomic identity was examined based on DNA barcoding using DNA sequences in the mitochondrial cytochrome c oxidase 1 (CO1) gene region and provided with the aid of available morphological keys and gene sequence information (Kondo, 2008; Lopes-lima et al., 2020). Total genomic DNA was extracted using a PureGene DNA isolation kit (Applied Biosystems, Waltham, MA, USA), following the manufacturer’s protocol. The extracted DNA was then amplified for the 658-bp CO1 region fragments by polymerase chain reaction (PCR) with universal primers LCO1490 and HCO2198 (Folmer et al., 1994) (Supplementary information 2).

The PCR products were purified using reagents with PEG-based purification procedure. The purified products were cycle sequenced using BigDye Terminator v1.1 or v3.1 (ABI: Applied Biosystems, Waltham, MA USA), and were sequenced on an Applied Biosystems 3130.

### Demographics survey

In October 2021, approximately 30 min were spent at two intensively monitored sites (M4 and M6), and relatively large adult Unionoidae sp. individuals were collected. Personnel (JN Negishi) with a dry suit hand-searched the bottom surface by one hand. Because small-sized juveniles were unlikely to be quantitatively collected by this method, smaller individuals were thoroughly hand-searched by spending another 30 min by combining two methods (i.e., collecting bed sediment with two hands and a hand net and examining the contents very carefully). A total of 262 individuals were collected (85 and 177 individuals in M4 and M6, respectively). Shell length was measured, followed by examination of the occurrence and level of gravidity by carefully prying the shell open to inspect the gills. Mussels were enclosed in a mesh net and kept in the middle of the collection area at the lake bottom (water depth of approximately 60 cm) until measurements were taken. Daytime dissolved oxygen was much lower in winter periods under ice than in summer according to the data previously measured; summer dissolved oxygen was much lower in M4 than in M6 (Supplementary information 3a). Also, fish communities were similar between two sites (Supplementary information 3b)

At an interval of one month to several months starting from October 15, 2020, the same sets of Unionidae sp. adult individuals (42 and 32 individuals in M4 and M6, respectively) were collected and checked for their survival and level of gravidity, as described above. In this step, we assigned them into one of nine categories (modified from Negishi et al., 2018): at a stage before gravidity or no reproduction (non-gravid); at an early stage of gravidity (early-gravid) having slightly swollen cream-colored demibranchs; fully gravid (egg-gravid) with fully swollen cream-colored demibranchs; with glochidia (grochidia-gravid) having fully swollen amber-colored demibranchs; with ripe glochidia (ripe-grochidia-gravid); having partially flattened (<10%) amber-colored demibranchs (early-released); flattened (<50%) amber-colored demibranchs (late-released); flattened (<90%) amber-colored demibranchs (very-late-released); and having slightly swollen demibranchs without glochidia (spent). The dead individuals were recorded and removed when they were found. Mussels were collected through 15-cm-diameter holes drilled through the ice layer during winter.

Other individuals (122 adults in M4, 19 juveniles, and 118 adults in M6) were used for annual measurements of their survival and observations of gravidity cycles and sex in parallel experiments. Approximately half of the mussels from M6 were used for parallel experiments (two weeks of rearing and measurements for their metabolic activities in the laboratory and immediate release back into the field; Wu et al., 2023) but treated together with the rest of the mussels without experiments because preliminary examinations of the effects of this treatment did not show significant differences in comparison with the remaining mussels in M6 in terms of mortality; other individuals were also used as a control to check whether monthly monitoring caused an increased/decreased rate of mortality as an artifact). In November 2021, mussels were again measured for their survival and gravidity levels in correspondence with the records of their gravidity in October 2020. All individuals were kept in mesh bags throughout the study period, according to gravidity and size classes, in such a way that their gravidity status in October 2020 could be tracked to November 2021 to understand the gravidity cycle at an annual scale.

### Analyses

The obtained DNA sequences were combined with those of closely related species sequences in a database (NCBI). Closely related species sequences were found in the BLAST search, and those belonging to bivalves with >90% identity to the searched individual (BI046 in Supplementary information 1) were downloaded (in total 40 sequences; Supplementary information 4). These sequences were aligned and truncated at maximum common base-pair lengths (602bp) and used to construct a phylogenetic tree based on the neighbor-joining (NJ) method with a rapid bootstrapping procedure using 1,000 bootstrap analyses using MEGA X (Kumar et al., 2018).

The mussel mortality rate was calculated as the proportion of surviving individuals in relation to the initial abundance at the beginning of October 15, 2021 (%). For the annual measurements of mortality rates, the mortality rate was separately determined according to gravidity (gravid and non-gravid) and size class (adults and juveniles). Chi-square tests were also performed to compare the mortality rates among sizeLgravidity groups between the two sites, as well as within each site. All analyses were conducted using R (R Core Team, 2022) and relevant packages, such as “ggtree” and “ggplot2” (Yu et al., 2017). Statistical significance (α) was set at p =0.05.

## Results

### Species identification

The same species identity of the collected Unionidae sp. individuals including those at two intensively monitored sites was well supported based on the high divergence of COI compared with other closely related other species in the genus of *Buldowskia* (*B. kamiyai* and *B. shadini*) (Fig. 2). The sequences of our own Unionidae sp. and other reported sequences of *B. iwakawai* in NCBI database (in total of 59 sequences) were all included in the clade branched off from the one that belonging to *B. kamiyai*. *Anemina arcaeformis* was substantially diverged from *Buldowskia* species.

**Figure 2.**
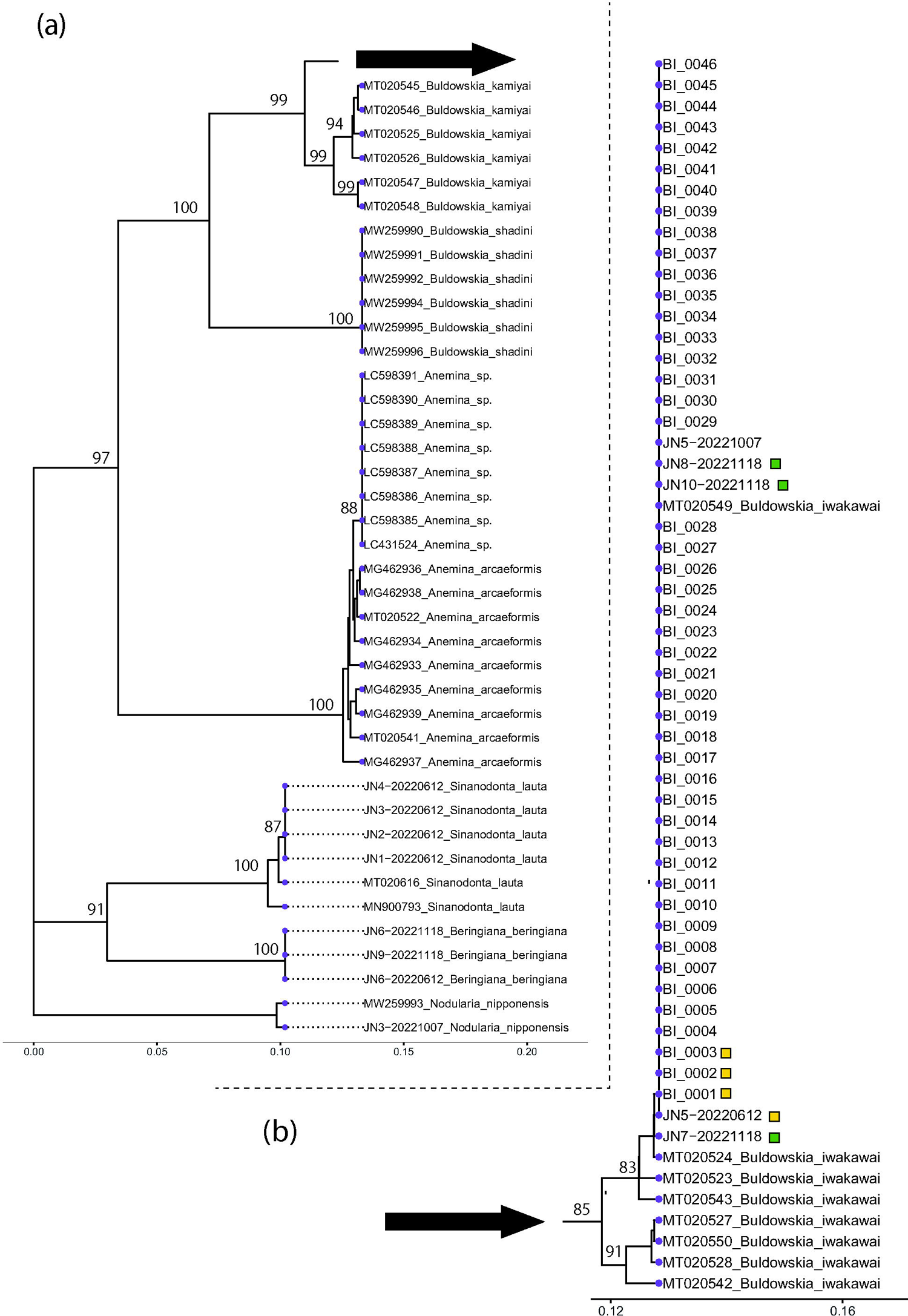
Phylogenetic tree showing the genetic position of the studied Unionoid sp. in relation to other species in the family Unionidae found in the Ishikari River and sequences collected from the NCBI database. Those accompanied by yellow and green boxes are individuals collected from sites M6 and M4, respectively. Bootstrap values are also shown for those with >80.

Only two haplotypes with one mutation site among 59 individuals collected from 11 floodplain lakes, including two lakes intensively monitored. One haplotype was identical to *B. iwakawai* in NCBI database (MT020549) and the other also close to haplotypes of *B. iwakawai* in the database, confirming our samples as *B. iwakawai*. *Buldowskia iwakawai* showed two clades, one found in Korea and western Japan (MT020527, MT020550, MT020528, MT020542) and another in central and eastern Japan (MT020523, MT020543, and T020524) Japan (Supplementary information 1). Our samples in Hokkaido were all included in the later clade.

### Population demographics

The target Unionidae sp. in the two intensively monitored sites had contrasting demographics in terms of size distribution. At site M6, there were abundant juvenile mussels, including newly recruited mussels with shell sizes <3 cm, whereas no such individuals were caught in an additional juvenile search at site M4 (Fig. 3). Gravid mussels in M6 contained relatively large gravid individuals compared to sympatric non-gravid mussels, whereas the size range of the mussels was similar between the non-gravid and gravid groups in M4.

**Figure 3.**
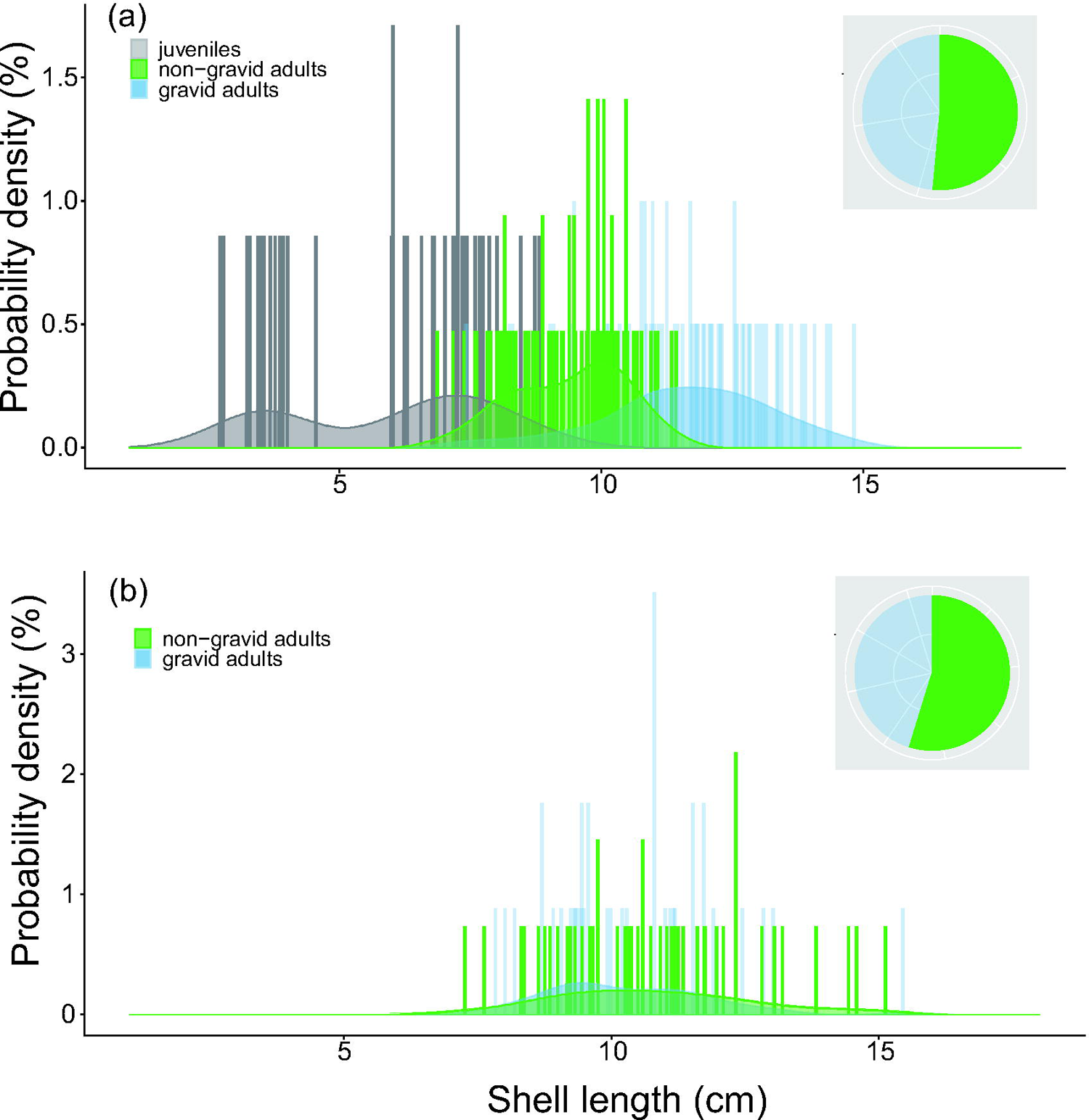
Size distribution of the studied Unionoidae sp. in relation to their size class and gravidity at sites M6 (a) and M4 (b), measured in October 2021. The pie chart in each panel shows the proportion of gravid and non-gravid mussels.

The gravidity levels of Unionidae sp. gradually progressed over time, and reproduction mostly ceased by June at both sites (Fig. 4). Site M4 was not measured in January and February due to heavy snow cover (>1 m depth) on the surface ice. The continuous observations at site M6 and observations of gravidity stage changes between December and April at site M4 suggest a peak of glochidia release in March, April, and May. Mortality was low in the early occasions, gradually increased until February (at least in site M6), and reached the highest mortality rates in April (the end of winter periods) at approximately 40% in both cases (38.1% in M4, and 31.3% in M6).

**Figure 4.**
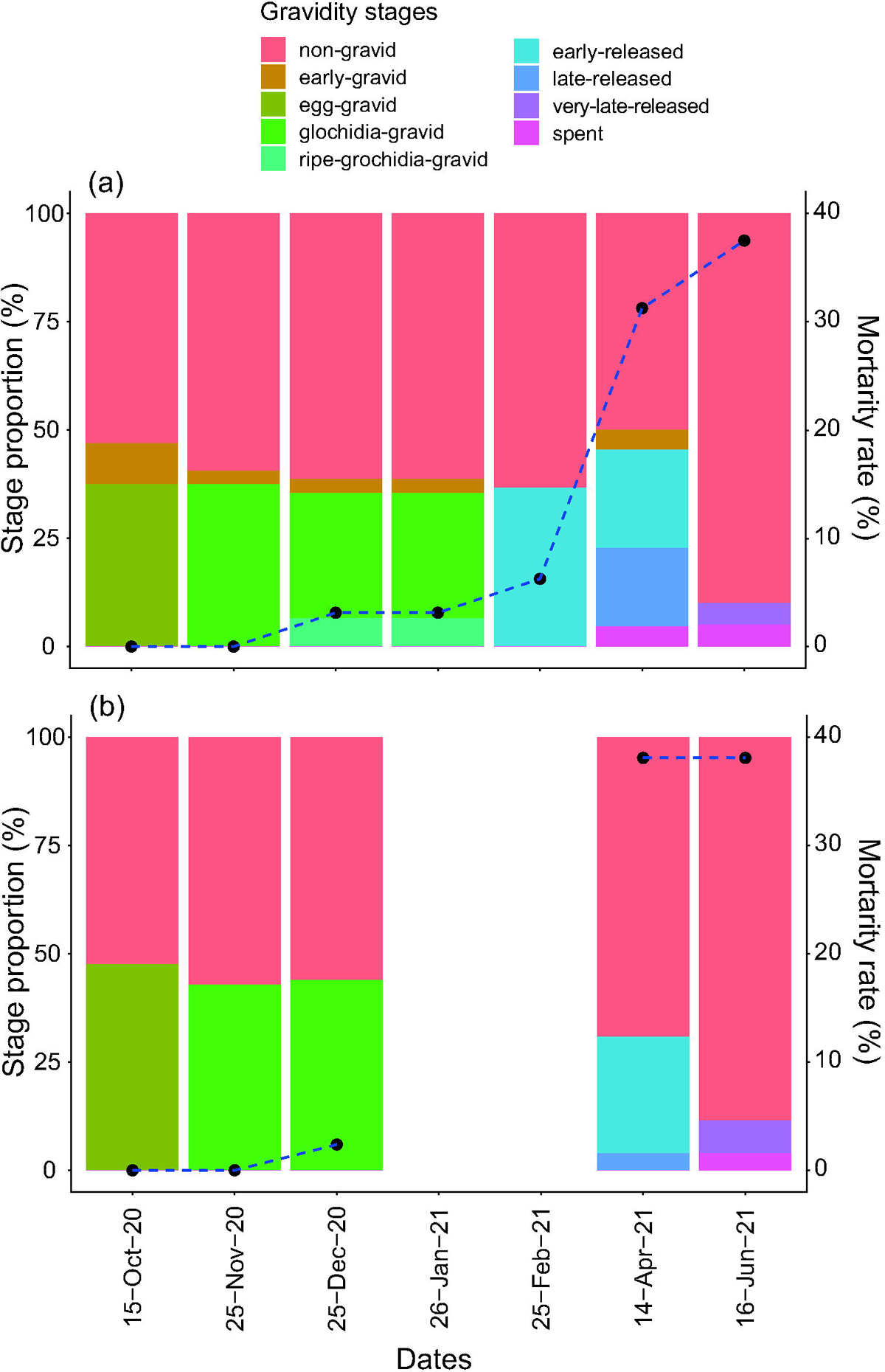
Gravidity stage progression of the studied Unionoidae sp. in relation to their mortality since the onset of observations in October 2020 at the site M6 (a) and M4 (b). Data in January and February at the site M4 were not shown due to data unavailability as a result of heavy snow depth and inaccessibility to the site.

Annual mortality rates differed significantly between sites only for gravid mussels, with the rate being much higher in M4 when compared between the same groups at two sites (p<0.01), and differed among groups in M4 (p<0.01) (Fig. 5). The mortality rate of gravid adults (ca.75.0%) was much higher than that of non-gravid mussels in M4 (ca.45.2%); thus, an approximately 30% mortality increase was observed after the winter monitoring period (non-winter months) in M4. As a result, the mortality of gravid mussels was higher than that of non-gravid mussels in non-winter months by 2.7 times (36.9% and 13.9%). In contrast, the annual mortality of other adult groups did not differ much from those observed in winter months. The mortality of juveniles was the lowest among all groups, although the juveniles were not statistically distinguishable from the two other adult groups in M6. Except for one gravid individual in M4, which did not become gravid in the second year, gravid/non-gravid categorization did not change based on the assessment after one year; other surviving gravid mussels were persistently gravid in two years.

**Figure 5.**
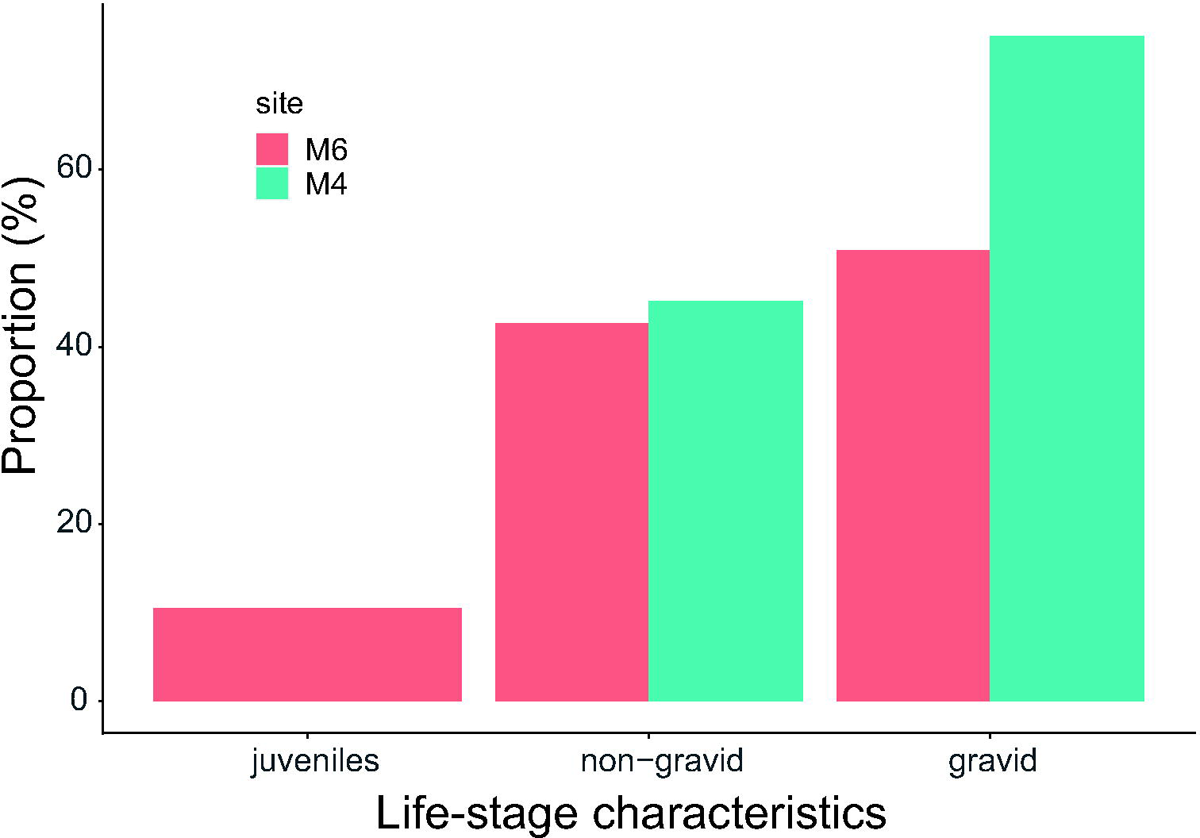
Annual mortality of the studied Unionoid sp. according to life-stage characteristics (size and gravidity determined in October 2020). Data of juveniles in the site M4 was not shown because no juveniles were found.

## Discussion

Harsh and long winters pose considerable challenges in field observations of ecosystems, partially explaining the underrepresented roles of winter in material cycles, native and non-native distributions, and population dynamics of organisms (Campbell et al., 2005; Huusko et al., 2007; Koizumi et al., 2017; Miura et al., 2017). The need for science-based implications for conservation and habitat management of imperiled taxa is not limited by these challenges (Negishi et al., 2018; Miura et al., 2023a, b). This study sheds new light on the winter-mediated mortality of freshwater mussels in shallow eutrophic floodplain lakes. Our prediction was partially supported because mussel mortality was higher in winter periods than non-winter periods. However, the second prediction, that non-winter mortality is more severe in the shrinking population, was not supported, as the mortality level of mussels in winter was comparable between the two sites. The causes of apparently impaired reproduction in M4 need to be further narrowed to more complicated processes.

Genetic barcoding analyses confirmed that the targeted species was *B. iwakawai*; thus, we conclusively revised the identity of previously reported mussels, *A. arcaeformis*, in the areas of Negishi et al. (2018) and Izumi et al. (2020). Recent studies have reported the occurrence of *Buldowskia* species, such as *B. shadini* and *Anemina* species in China (native habitat range) as well as confined areas in Japan (Kawase et al., 2021; Uchino et al., 2021), suggesting localized non-native species invasions in Japan. Based on CO1 sequence divergences among individuals, *B. iwakawai* in Hokkaido and central Japan was distinguished from others reported in western Japan and Korea, which is in line with the spatial geographical divergence of this species without reasonable signs of artificial introductions in its non-native areas. Thus, the current findings support the notion that *B. iwakawai* in the Ishikari floodplain is likely within the natural distribution of this species, with no invasion of *A. arcaeformis* and other Buldowskia species. This clearly underscores the need for conservation and improvement of the *B. iwakawai* habitat in the area given its declining population and MLIT’s policy in environmental restoration of the area. DNA-based species identification is a powerful and promising tool to monitor the invasion of mussel species, population dynamics, and distribution of native species and their hosts, as shown in other studies (Zieritz et al., 2012; Marshall et al., 2018). Our findings also support the notion that improved morphological keys combined with genetic information are preferable for improving mussel conservation and habitat management from a cost-benefit perspective. (Miura et al., 2019; Wu et al., 2023).

The glochidia release timing of unionid species has been reported in other species, such as margaritiferids and unionoids (Österling, 2015; Haggerty & Garner, 2000), including some winter-breeders (Fukuhara et al., 1987; Kondo et al., 1987; Roberts & Barnhart, 1999; Jones et al., 2010). For *B. iwakawai*, the timing of glochidia release was inferred to occur sometime in spring based on laboratory experiments (Negishi et al., 2018). Our results are unique in that the temporal progression of mussel mortality was revealed in relation to the gravidity stages. The results clearly showed that winter mortality was high when the timing was highly synchronized with that of glochidia release. This winter-period mortality is unlikely to be an artifact of repeated examinations of the stage because the control group mussels, which were untouched between measurement occasions, did not show systematically reduced mortality. We believe that the cause of mortality was related to decreased dissolved oxygen under the ice and the physiological stresses caused by it, as commonly reported in productive shallow lakes (Magnusonl et al., 1985; Hilt et al., 2015). No continuous dissolved oxygen data were available; however, spot measurements of daytime DO support this argument. Because high winter mortality is common for both gravid and non-gravid mussels, this pattern is not driven by gravidity. Instead, it is inferred that physiological stress levels for mussels reached a certain threshold, then the oxygen depletion was stable over time or oxygen depletion level gradually increased around this time of the year. Further detailed monitoring of temporal changes in oxygen levels under ice is necessary to address these hypotheses.

Juvenile mortality in winter was much lower than that in adult mussels, as indicated by the remarkably low annual mortality of approximately half of adult mortality at the same site. In a parallel study, juvenile mussels were found to be significantly less metabolically active than adult mussels (Wu et al., 2023), suggesting that size-dependent mortality was manifested by a combination of seasonally intensified oxygen depletion and size-specific metabolic rates of mussels (i.e., physiological demands). With the fact that glochidia release per se was not differently limited in two sites together with the low mortality of juveniles in winter periods, bottlenecking life stage for the impaired population limitations was narrowed down to the stage between released glochidia and the very early juvenile stage before reaching a size of 3 cm in shell size. Two mechanisms, host-fish limitation and habitat condition of newly born juveniles (< 3 cm in shell size), are the most likely causes, as shown in previous studies (Stoeckl et al., 2015). Host-fish limitations in summer are unlikely because previous surveys revealed that both sites were similarly abundant and rich in fish communities, including the regionally important fish host, Chestnut goby (*Gymnogobius castaneus*) (Supplementary information 3b; Negishi et al., 2018).

Non-winter mortality (i.e., annual mortality rates excluding the contribution of winter mortality) was informative of the potential cause of contrasting population structures. Selectively imposed mortality on gravid mussels in the non-winter period was the most apparent difference between populations. Spot measurements of DO in summer suggested more severe hypoxic conditions and intensified physiological stresses on mussels in M4 because metabolic rates can increase with temperature (e.g., Haney et al., 2020). This was partially driven by the morphological differences between the two lakes. M4 is shallower than M6, and thus the macrophyte cover on the lake surface in summer was different from M4 (> 80 %) and M6 (< 10% covered), causing a higher frequency and intensity of respiration-derived anoxia in summer (Negishi et al., 2014). These harsh hypoxic environments could disproportionately affect newly born juveniles because young post-glochidial mussels are highly sensitive (Dimock & Wright, 1994). However, it is unclear why gravid mussels are selectively and more severely affected. Although metabolic rates were not measured during the summer, it is unlikely that gravid (female) mussels were characterized by higher metabolic rates because no glochidia were held at that time. One plausible explanation is the delayed carry-over effects of physiological stress during winter breeding periods. Physiological costs associated with glochidia development and release, which are further exacerbated by winter low DO, might make gravid (female) adult mussels more vulnerable to environmental stresses in subsequent warmer non-winter periods (Johns et al., 2010). This selective loss of gravid mussels may be the cause of the slight preference of non-gravid mussels in terms of demographics. Larger mussels were more abundant in the well-reproducing lake, which is also consistent with the extended longevity of gravid mussels under benign summer habitat conditions. Thus, in addition to the presence of juveniles, the unbiased composition of gravid to non-gravid mussels and the presence of large-sized gravid mussels could be useful indicators of healthy populations. The rare occurrence of hermaphrodites and the continuous annual reproduction of gravid mussels found in this study support this notion.

### Implications for conservation

In conclusion, winterkill does not explain the contrasting demographics of winter-breeding *B. iwakawai* populations in the Ishikari River floodplain. The winterkill of *B. iwakawai* in the studied life cycle stages was not a strong independent driving factor for the troubled population of the study species. Instead, winterkill is likely to come into play as a significant bottleneck when combined with summer environmental conditions that are critically stressful, owing to low dissolved oxygen levels. Under such conditions, a chain of adult abundance decreases in winter, high mortality of gravid mussels and newly born juveniles occurs in summer, and population size decreases with a lack of juveniles at annual or longer timescales. Recent studies have indicated that multiple stressors can limit the community structure, distribution, and reproduction of freshwater species, including freshwater mussels, in complex ways (Birk et al., 2020; Miura et al., 2023a). Our findings extend the multiple-factor paradigm and underscore the importance of considering survival limitation factors acting at multiple life stages of freshwater mussels at different times of the year. In this context, demographic information of the population provides crucial knowledge for future conservation research that attempts to elucidate the mechanisms behind and decelerate the population decline trends of freshwater mussels. In this context, demographic information of the population provides crucial knowledge for future conservation research that attempts to elucidate the mechanisms behind and decelerate the population decline trends of freshwater mussels.

DNA-based evidence supports the native origin of this species and proposes that this functionally important taxon can act as an aquatic indicator species in future floodplain restoration projects in the Ishikari River, as in other degraded floodplains (Negishi et al., 2013; Nagayama et al., 2023). Furthermore, ice-cover effects in lakes are not limited to parasite-host relationships and can be extended to food-web interactions (Hilt et al., 2015), highlighting the need for further studies on this topic in warming climates. From the perspective of conservation and management of the habitats of this native species, improvements in summer conditions are necessary, while attention should be paid to seasonal contexts, especially in winter. Increases in water circulation rates and the alleviation of hypoxic conditions are options for short-term habitat improvement. The effectiveness of this approach and the validity of our conclusions can be tested by diverting well-oxygenated water into concerned waterbodies, followed by temporal monitoring of demographic changes. In the long run, a more process-oriented approach that can lead to reduced intensity and frequency of hypoxia formation, including treatment of incoming sediment and overgrowth of aquatic plants and their deposition, needs to be considered. In the current planning of habitat restoration in the Ishikari River floodplain, several degraded floodplain lakes will be selected to conduct pilot experimental projects that provide empirical evidence to promote the full-scale restoration of degraded floodplain ecosystems. Our findings facilitate the generation of hypotheses, predictions, and prioritized site selection with monitoring schemes covering harsh winter periods.

## Author contributions

N Negishi: Conceptualization (equal), data collection (equal), data curation (equal), formal analysis (equal), funding acquisition (equal), and leading writing. H Izumi: Conceptualization (equal), data collection (equal), formal analysis (equal), and data curation (equal). J Wu: Data collection (equal), formal analysis (equal), and data curation (equal). S Fukui: formal analysis (equal). I Koizumi: Conceptualization (equal); funding acquisition (equal). All the authors contributed critically to the drafts and approved the final manuscript for publication.

## Acknowledgements

We are grateful to the Sapporo Regional Office of Hokkaido Development Bureau, MLIT for data collection and field assistance. This study was partly supported by the research fund for the Tokachi and Ishikari rivers provided by MLIT (18056588) and JSPS KAKENHI (22H03787).

## Ethical approval

No ethical violation was occurred in this research.

## Data availability statement

All the data used in the paper will be deposited in Dryad and DNA data bank of Japan

