## Supplementary material for "Does winterkill explain contrasting demographics of winter-breeding freshwater mussel populations in Ishikari River floodplain?": SM1

**Supplementary information 2**

PCR assay (a) and conditions (b) in the present study.

(a)

Total reaction volume: 25μl

PCR premix: GoTaq PCR buffer (Promega) or DreamTaq Hot Start PCR Master Mix (Thermo Scientific): 12.5μl

Primers (10 μM): 0.5 μl each for forward and reverse primers

Distilled water: 10.5 μl

Template total genome extract: 1 μl

(b)

| Steps | Temperature (℃) | Duration (min) | Cycle repeat | Cycle times |
| --- | --- | --- | --- | --- |
| Pre-heating | 95 | 2 |  |  |
| Denaturing | 94 | 1 | Yes | ×25 cycles |
| Annealing | 45.5* | 0.5** | Yes |  |
| Extending | 72 | 1 | Yes |  |
| Final extending | 72 | 5 |  |  |
| Storing | 10 | ∞ |  |  |

*In some cases, annealing temperature was increased or decreased within a range of 2 ℃ to obtain the best PCR amplification efficiency.

** In some cases, annealing duration was increased to 1 min to obtain the best PCR amplification efficiency.
