## Supplementary material for "Does winterkill explain contrasting demographics of winter-breeding freshwater mussel populations in Ishikari River floodplain?": SM2

**Supplementary information 3**

Dissolved oxygen measured in sites M4 and M6 (A). Own data was measured in February 2022 in both lakes whereas MLIT data was measured as a part of government assessments of lakes in August and October 2006. All data was collected at the baseflow conditions without large floods within two weeks and ice cover (August and October). Fish survey was conducted in August 2016 by similar CPUE efforts using seine nets, electro-fisher, box trap, and cast nets (B); the number of individuals caught was shown.

(A)

| Data type | Site | Measurement timing | DO (mg/L) |
| --- | --- | --- | --- |
| Own | M4 | February 2022 | 4.1 |
| Own | M6 | February 2022 | 3.8 |
| Own | M4 | July 2022 | 5.7 |
| Own | M6 | July 2022 | 6.9 |
| MLIT | M4 | August 2006 | 3.3 |
| MLIT | M6 | August 2006 | 7.5 |
| MLIT | M4 | October 2006 | 10.7 |
| MLIT | M6 | October 2006 | 10.6 |

(B)

| Taxonomic names | M4 | M6 |
| --- | --- | --- |
| *Gymnogobius castaneus* | 179 | 183 |
| *Pseudorasbora parva* | 53 | 63 |
| *Hypomesus olidus* | 0 | 83 |
| *Pungitius sinensis* | 71 | 1 |
| *Rhodeus ocellatus* | 60 | 4 |
| *Carassius buergeri* | 14 | 41 |
| *Rhynchocypris percnurus sachalinensis* | 2 | 14 |
| *Carassius* sp. | 12 | 2 |
| *Tribolodon* sp. | 7 | 0 |
| *Tribolodon sachalinensis* | 1 | 0 |
