## Supplementary material for "Does winterkill explain contrasting demographics of winter-breeding freshwater mussel populations in Ishikari River floodplain?": SM3

**Supplementary information 4**

Specifications of CO1 sequences of closely related unionid mussels. Sample ids are accession numbers in NCBI GenBank database.

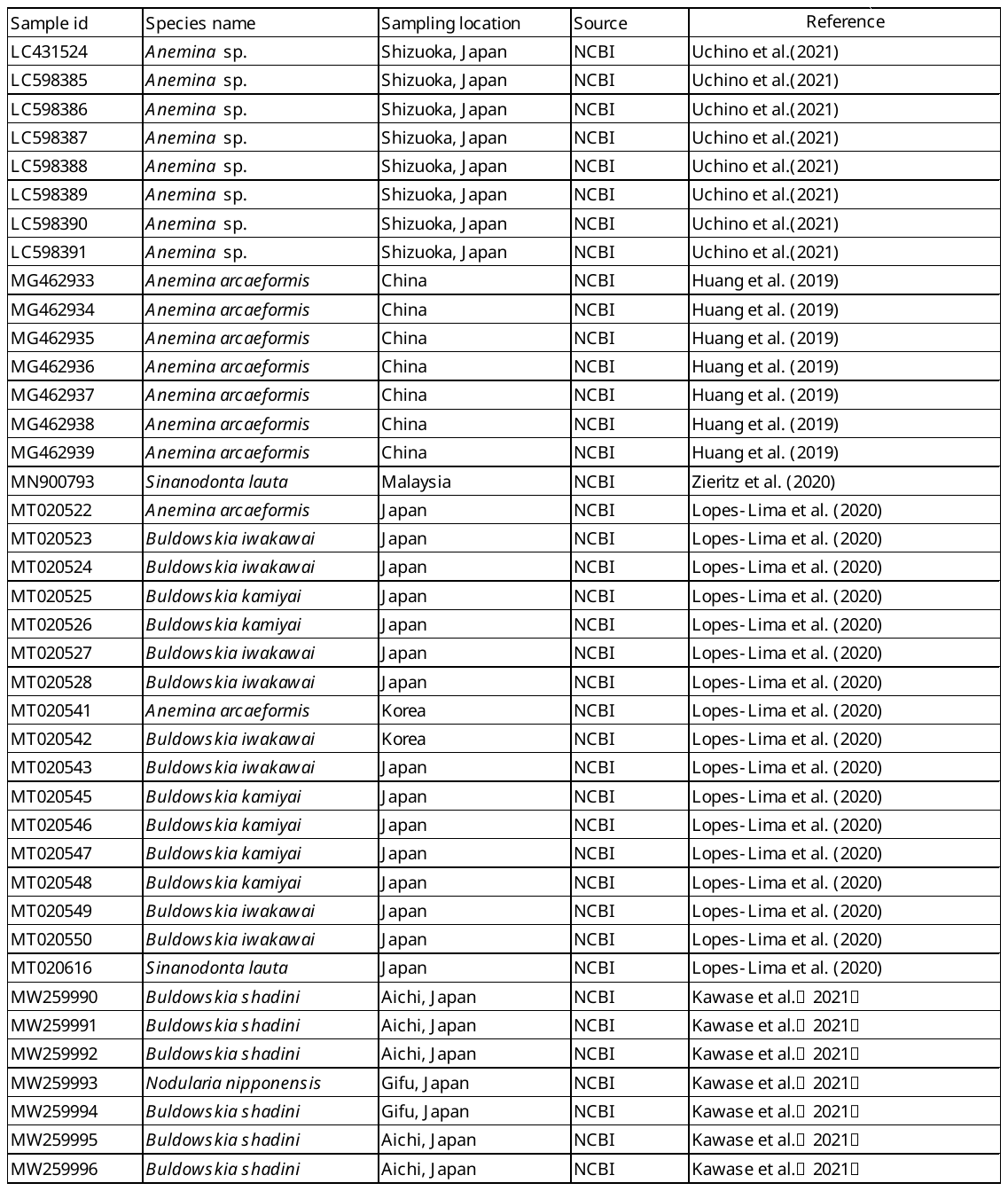
