## Supplementary material for "Does winterkill explain contrasting demographics of winter-breeding freshwater mussel populations in Ishikari River floodplain?": SM4

**Supplementary information 1**

Specifications of collected unionid mussels for CO1 sequences in this study.

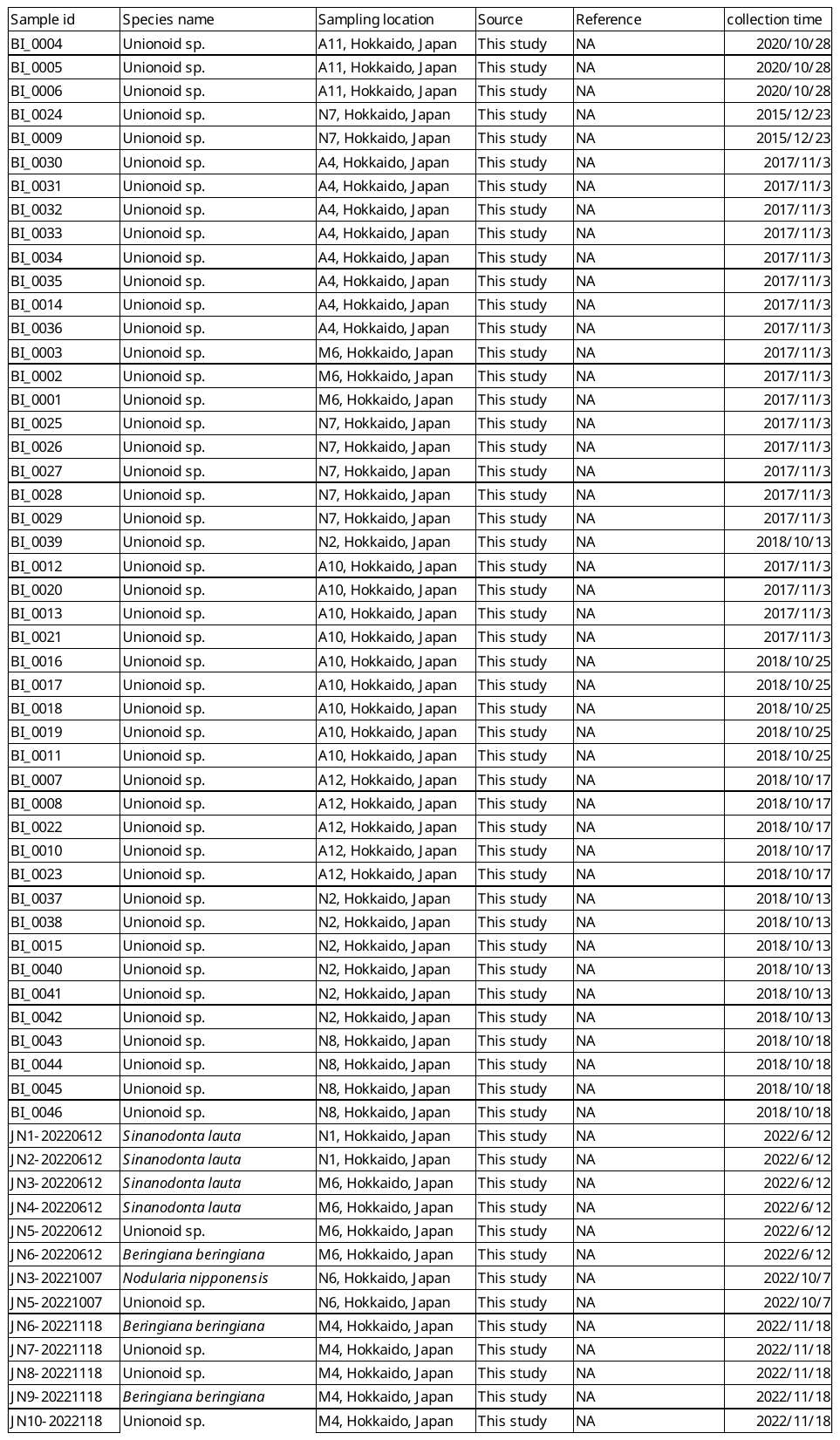
